# Influence of Sodium Salicylate on Adventitious Organogenesis of a Commercial Cucumber Cultivar

**DOI:** 10.1101/2022.09.16.508332

**Authors:** Jorge Fonseca Miguel

## Abstract

The effect of different concentrations of sodium salicylate (NaSA), a phenolic compound, on *in vitro* callus and shoot regeneration of cucumber (*Cucumis sativus* L.) was investigated. Four-day-old cotyledon explants from the Spanish cultivar ‘Marketer’ were employed. MS-derived shoot induction medium containing 0.5 mg L-1 IAA and 2.5 mg L-1 BAP was supplemented with NaSA. Frequency and shoot number were enhanced by 1.5-fold with NaSA at 0.1 μM. Higher salicylate levels led to increased callus formation and decreased shoot regeneration. The application of sodium salicylate at a specific concentration showed a positive trend in *in vitro* adventitious organogenesis of a commercial cucumber cultivar. Some probable mechanisms that may underlie the beneficial effects of salicylic acid/salicylates on *in vitro* regeneration were also discussed.

## 1. Introduction

Sodium salicylate (sodium 2-hydroxybenzoate; NaSA) is the sodium salt of salicylic acid (SA) (PubChem, 2022a). SA (2-hydroxybenzoic acid) is an endogenous growth regulator and a signaling molecule involved in the regulation of various physiological processes related to plant growth and development, as well as in the modulation of plant responses to different abiotic and biotic stresses (Mohamed et al. 2020; PubChem, 2022b).

Exogenous application of SA may affect a variety of plant processes and attributes, such as photosynthesis and photosynthetic pigments (Khan et al. 2003; Maurya et al. 2019), respiratory pathways (Khan et al. 2003), ethylene biosynthesis (Khan et al. 2013), seed germination (Lee et al. 2010), flower induction (Cleland and Tanaka 1979), thermogenesis (Rhoads and McIntosh 1992), ion uptake and transport (Harper and Balke 1981; Gondor et al. 2016), nitrate reductase activity (Fariduddin et al. 2003), protein content (Latif et al. 2016), growth parameters (Damalas 2019), biomass production (Maurya et al. 2019), plant water relations (Hayat et al. 2010), stomatal closure (Khan et al. 2003), accumulation of osmolytes (e.g., proline, glycine betaine, polyamines, and soluble sugars) (Khan et al. 2014; Madany et al. 2020; Shemi et al. 2021), secondary metabolite content (e.g., alkaloids, flavonoids, and phenolics) (for review see Nandy et al. 2021), allelopathic properties (Shettel and Balke 1983), antioxidant defense system (Maurya et al. 2019), and senescence (Rivas-San Vicente and Plasencia 2011). It can also alleviate environmental stresses, such as heat (Khan et al. 2013), cold (Saleem et al. 2020), drought, (Sohag et al. 2020), UV-B/C radiation (Li et al. 2014; Abrun et al. 2016), salinity (Elazab and Youssef 2017), and heavy metals toxicity (Sharma et al. 2020), and protect plants from a range of pathogens (Bakker et al. 2014). The effect of exogenous SA is influenced by factors such as the plant species, its stage of development, tissue and organ type, mode of application, dose and duration of exposure, environmental and culture conditions, and its endogenous level in the plant (Horváth et al. 2007; Janda et al. 2014; Kračun-Kolarević et al. 2015; Bhutia et al. 2018). SA application has been reported to modulate plant functions in a dose-dependent manner (Maurya et al. 2019). Inadequate levels of SA can also lead to oxidative stress (Gondor et al. 2016). SA-like effects have been reported with structurally similar compounds, such as NaSA and ASA (acetylsalicylic acid). Some authors have previously described improved *in vitro* plant regeneration with exogenous SA and ASA. This is investigated in the present work using NaSA.

Cucumber (*Cucumis sativus* L.) is an economically important species belonging to the Cucurbitaceae family. Its world production, including gherkins, ranked third among vegetable crops in 2020, with 91,258,272 tons (FAOSTAT, *http://www.fao.org/faostat/en/#data/QC*). This species is an important source of valuable nutrients and bioactive compounds, used not only as food but also in cosmetics, therapeutics, and perfumery (Uzodike and Onuoha 2009; Uthpala et al. 2020).

*In vitro* regeneration in cucumber has been established and can be achieved through organogenesis and somatic embryogenesis. The most frequently used explants are cotyledons (Chee 1990; Selvaraj et al. 2007; Miguel 2017), hypocotyls (Ziv and Gadasi 1986; Chee 1990; Grozeva and Velkov 2014), and leaves (Malepszy and Nadolska-Orczyk 1983; Lou and Kako 1994; Seo et al. 2000). Regeneration from protoplasts (Jia et al. 1986; Trulson and Shahin 1986; Burza and Malepszy 1995) and suspension cultures has also been reported (Malepszy and Solarek 1986; Bergervoet et al. 1989; Kreuger et al. 1996). However, regeneration in cucumber is still not optimal (Wang et al. 2015) and highly dependent on the genotype (Wehner 1981; Punja et al. 1990; Miguel 2021a, b). An efficient and reproducible regeneration system is fundamental to address different biotechnological approaches, such as secondary metabolite production, germplasm conservation, and large-scale multiplication in this species.

This study aimed to evaluate the influence of different concentrations of sodium salicylate on *in vitro* adventitious organogenesis of a Spanish commercial cucumber cultivar.

## 2. Materials and Methods

### 2.1. Plant material and in vitro regeneration

Cucumber seeds of the Spanish commercial cultivar ‘Marketer’ (Semillas Fitó S.A.) were the starting material. Obtaining axenic explants and *in vitro* regeneration followed the methodology previously described by Miguel (2021a). Four-day-old cotyledons with proximal and distal ends excised by 1-2 mm deep were used as explants. They were cultivated for 3 weeks on MS-derived shoot induction medium containing 0.5 mg L-1 IAA and 2.5 mg L-1 BAP and supplemented with sodium salicylate (0.0, 0.1, 1.0, and 10.0 μM; Sigma–Aldrich, St. Louis, MO, USA). Callus frequency (CF) and callus extension index (CEI) were then determined. The CF (mean±SE) is the proportion of explants with callus on the explant cut surface, and the CEI (mean±SE) corresponds to the extent of callus on the cut surface, and was graded as follows: 0 = absence of callus; 1= trace of callus; 2 = callus on less than half; 3 = callus on half or more; 4 = callus covering the entire extent. Organogenic structures where then culture for 2 weeks on shoot development and elongation medium containing 0.2 mg L-1 KIN. At the end, an evaluation of shoot regeneration frequency (SRF) and shoot number index (SNI) was made. SRF (mean±SE) is the proportion of explants forming shoots, and SNI (mean±SE) was scored according to the number of shoots per explant as follows: 0 = absence; 1 = one shoot; 2 = two shoots; 3 = three or more shoots. Individualized shoots were then rooted on hormone-free MS medium and, within 3 to 4 weeks, the plantlets were ready for acclimatization (data not shown). *In vitro* culture media were solidified with 0.8% (w/v) agar and adjusted to pH 5.7 before autoclaving. Incubation was performed in a culture chamber at 26 ± 2 ºC under a 16L:8D h photoperiod. Light was provided by cool-white fluorescent lamps (Grolux, Sylvania) at a photon fluence rate of 90 μmol m–2 s-1.

### 2.2. Data analysis

The experiment was a single factor design with four levels of sodium salicylate (NaSA) arranged in a completely randomized design. Twelve replicate flasks with six explants each were used in each treatment, except for 10.0 μM NaSA with six replicate flasks. The R statistical computing environment (version 4.0.4; R Core Team 2021) was used to perform all statistical analyses. Non-linear regression analyses were applied to the data to compare treatment means. The R packages ‘stats’ (R Core Team 2021), ‘COMPoissonReg’ (Sellers et al. 2019), and ‘pscl’ (Jackman 2020) were used to conduct logistic regression, COM-Poisson regression, and Hurdle regression, respectively. The goodness of fit of the models was assessed using the Akaike Information Criterion (AIC; Akaike 1973) and the Bayesian Information Criterion (BIC; Schwarz 1978). The significance level was set at P ≤ 0.06 for shoot variables and at P < 0.05 for all other analyzes.

## 3. Results

### 3.1. Frequency and extension of callus

Results are presented as mean ± SEM (Table 1), with significance determined as P < 0.05. Callus formation occurred within the first two weeks of culture, starting at the cut ends of the cotyledon explant and scoring 100% frequency. The application of 0.1 μM sodium salicylate (NaSA) produced the lowest callus extension (1.22±0.05), with significant differences from the other treatments that did not differ from each other. Callus extension scores lie between the arbitrary values of 1: traces of callus, and 2: callus in less than half, at explant cut ends.

**Table 1.** Effect of adding sodium salicylate to MS-derived shoot induction medium containing 0.5 mg L-1 IAA and 2.5 mg L-1 BAP on *in vitro* callus and shoot regeneration from cotyledon explants of a commercial cultivar of *Cucumis sativus* L. Data are presented as mean ± standard error of the mean (SEM).

| Sodium salicylate |  | Callus regeneration frequency (%)<sup>&</sup> | Callus extension index<sup>¥</sup> | Shoot regeneration frequency (%)<sup>‡</sup> | Shoot number index<sup>§</sup> |
| --- | --- | --- | --- | --- | --- |
| $\mu\text{M}$ | $\mu\text{g L}^{-1}$ | | | | |
| 0.0 | 0 | 100 | $1.46 \pm 0.06^a$ | $30.56 \pm 5.47^b$ | $0.26 \pm 0.05^b$ |
| 0.1 | 16 | 100 | $1.22 \pm 0.05^b$ | $45.83 \pm 5.91^a$ | $0.38 \pm 0.06^a$ |
| 1.0 | 160 | 100 | $1.54 \pm 0.06^a$ | $23.61 \pm 5.04^b$ | $0.23 \pm 0.06^b$ |
| 10.0 | 1601 | 100 | $1.43 \pm 0.08^a$ | $20.00 \pm 6.86^b$ | $0.14 \pm 0.05^b$ |

<sup>&</sup>,<sup>¥</sup>Data were obtained after 3 weeks of culture on shoot induction medium.
<sup>‡</sup>,<sup>§</sup>Data were obtained after 2 weeks of culture on shoot development and elongation medium.
<sup>¥</sup>COM-Poisson regression, <sup>‡</sup>logistic regression, and <sup>§</sup>Hurdle regression were used to analyze the data. Mean values followed by different letters in the same column are significantly different: <sup>¥</sup> $P < 0.05$ ; <sup>‡</sup>,<sup>§</sup> $P \leq 0.06$ .

### 3.2. Frequency and number of shoots

Mean results ± SEM are shown in Table 1. Shoots formed from callus within 2-4 weeks of culture, mostly at the proximal end of the cotyledon explant. The highest frequency (45.83±5.91) and shoot index (0.38±0.06) corresponded to 0.1 μM NaSA, the lowest level, with differences compared to the NaSA-free control (P = 0.06) and the treatments with NaSA (P < 0.05), which did not differ from each other (P < 0.05). It represents a 1.5-fold increase compared to the control.

## 4. Discussion

The effect of sodium salicylate (NaSA) on *in vitro* regeneration of a commercial cucumber cultivar was studied.

NaSA is the sodium salt of salicylic acid (SA) (PubChem, 2022a) that dissociates to produce SA in solution (Palmer et al. 2019). SA is a growth regulator and a signaling molecule of phenolic nature ubiquitously distributed in plants (Raskin et al. 1990; Mohamed et al. 2020). Its basal levels vary greatly among species, even within the same family (Raskin et al. 1990; Rivas-San Vicente and Plasencia 2011). Exogenous application of SA and its derivatives (salicylates) may generate a range of metabolic and physiological responses in plants and affect their growth and development. The effect of some of these compounds on *in vitro* regeneration has been reported in various plant species.

Shoot regeneration of cucumber (cv. Poinsett 76) from cotyledon explants increased with 30 μM SA, whereas other levels at 10-50 μM SA were detrimental (Vasudevan et al. 2006). In cucumber (cv. Zhongnong 18), root formation using hypocotyl explants improved at specific SA levels (50 and 100 μM), not differing from the control (no SA) at 10 μM (Dong et al. 2020). In melon (*Cucumis melo* L.), callus and shoot formation from cotyledon explants increased with SA and ASA (acetylsalicylic acid) at 50-200 μM, while higher levels (300-1000 μM) had an inhibitory effect (Shetty et al. 1992). Shoot regeneration of *Hibiscus acetosella* and *H. moscheutos* from shoot meristem explants improved with 500 μM SA, while for 1000 μM SA, it did not differ from the control (no SA) (Sakhanokho and Kelley 2009). In potato (*Solanum tuberosum* L.), organogenesis from axillary bud explants may be promoted or inhibited by SA, ASA, or NaSA at 1 or 10 μM, depending on their concentration, the culture medium, and the hormonal balance. Somatic embryogenesis in callus culture of *Astragalus adsurgens* Pall. derived from hypocotyl segments increased with 75-200 μM SA, while 300 and 400 μM SA had no positive effect; SA levels did not affect callus growth, except for 400 μM SA, with a negative impact (Luo et al. 2001). Somatic embryogenesis of ten chir pines (*Pinus roxburghii* Sarg.) genotypes using shoot apical dome explants from mature trees was enhanced by applying SA at 7.2 μM (Malabadi et al. 2008). In suspension cultures of arabic coffee (*Coffea arabica* L.), SA at 10–6 μM had a positive effect on somatic embryogenesis (Quiroz-Figueroa et al. 2001). Based on the literature review, applied SA/salicylates act in a dose-dependent manner. It can be stimulatory, causing no relevant changes, or inhibit *in vitro* regeneration. The results of the present investigation are consistent with these findings.

In the current study, the frequency and extension of callus were not influenced by NaSA application, except for 0.1 μM NaSA with lower callus extension. The same NaSA concentration showed a positive trend in both shoot regeneration frequency and shoot number index, with a 1.5-fold increase compared to the control. In data not shown, higher levels of NaSA were studied. At 100 μM NaSA, both shoot regeneration frequency and shoot number index were similar to those at 10.0 μM NaSA. At 160 and 800 μM NaSA, callus formation decreased, and no shoots were formed.

The mechanisms underlying the effects of SA/salicylates on *in vitro* regeneration are complex and remain to be elucidated. Multiple factors and signaling pathways are involved.

Phytohormones are crucial for regulating physiological processes throughout the plant life cycle (Kosakivska et al., 2020). They play a key role in *in vitro* regeneration. Crosstalk of SA with other phytohormones has been described under different growth, development, and stress conditions (for reviews, see Rivas-San Vicente and Plasencia 2011; Koo et al. 2020; Hayat et al. 2021).

In the study by Torun et al. (2020), SA treatments changed endogenous levels of abscisic acid (ABA), cytokinins (CTKs), ethylene, IAA, and jasmonic acid, in leaves of barley (*Hordeum vulgare* L.) cultivars under control and saline conditions. Effects of SA depended on the treatment timing and the cultivar. The ability of SA to mitigate salt stress appears to result from increasing ROS scavenging and antioxidant enzyme activity, which is closely related to changes in endogenous phytohormones. In wheat, Shakirova et al. (2003) reported that exogenous SA changed the hormone content in ABA, IAA, and CTKs in seedlings during germination with rapidly reversible changes, increased cell division and root cell extension, plant growth, and yield. Under salinity, SA treatment lessened changes in phytohormones levels, and alleviated salinity damaging effects on seedling growth.

Auxins and cytokinins are two of the most critical factors for tissue culture (Szechyńska-Hebda et al. 2007). The balance between endogenous auxin and cytokinin signaling is crucial for *de novo* organ regeneration (Su and Zhang 2014, and references therein). In a recent study, SA treatments promoted adventitious root formation in cucumber explants through competitive inhibition of the auxin conjugation enzyme CsGH3.5 and increased the level of free IAA (Dong et al. 2020). GH3 family genes have been extensively identified and characterized in several plant species (reviewed by Liao et al. 2015), including cucumber (Wu et al. 2014). They may be responsible for cellular auxin homeostasis via the conjugation of auxin (mostly IAA) to amino acids (Liao et al. 2015, and references there in). The crosstalk between SA and auxin is also known to balance plant defense and growth (Zhong et al. 2021). Besides, SA and IAA share a common precursor (chorismate), the end product of the shikimate pathway (Pérez-Llorca et al. 2019). Little is known about the interaction between cytokinin and SA signaling pathways. Its role in *in vitro* regeneration remains to be explored. However, its involvement in other processes has been reported, but the mechanisms underlying its effects are complex. In *Arabidopsis thaliana*, it was suggested that cytokinin up-regulates plant immunity by elevating SA-dependent defense responses and in which SA, in turn, feedback inhibits cytokinin signaling (Argueso et al. 2012). Studies in japonica rice (*Oryza sativa* subsp. japonica) showed that cytokinins acted synergistically with SA to trigger defense gene expression (Jiang et al. 2013). In barley, SA treatments reduced the total endogenous cytokinin content in leaves of two of the three cultivars and increased it in the third. Under saline conditions (150 and 300 mM NaCl), SA treatments generally increased total CK levels, particularly at 300 mM salt (Torun et al. 2020). Therefore, if SA can interact with cytokinins, it is likely that it may also influence the *in vitro* regeneration process.

It is known that SA operates through many targets to mediate its many effects on plant processes (Klessig et al. 2016). A variety of SA-binding proteins (SABPs) have been identified in plants (Klessig et al. 2016; Pokotylo et al. 2019). Their wide range of affinities for SA, together with the varying SA levels in subcellular compartments, tissues, developmental stages, and following an environmental stimulus, provides great flexibility and involves multiple mechanisms by which SA can exert its effects (Klessig et al. 2016).

Plant tissue culture conditions can cause stress that affects the protocol performance. It may also lead to genetic and epigenetic alterations (Pacheco et al. 2008). Exogenous application of SA and salicylates may generate protective effects against a variety of biotic and abiotic stresses. Citrus exposed to NaSA in laboratory and field experiments alleviated heat, cold, and disease stresses (Mann et al. 2011). Applied NaSA attenuated the adverse effects of salt stress on wheat *(Triticum aestivum* L.) and stimulated growth by increasing photosynthetic rate and retarding dark respiration (Al-Hakimi 2001). The toxicity of the heavy metals Pb and Cu in duckweed (*Lemna gibba* L.) and cadmium (Cd) in maize (*Zea mays* L.) were reduced by exogenous NaSA (Duman et al. 2010; Gondor et al. 2016).

Stresses generated by *in vitro* culture, such as those resulting from excess ethylene, reactive oxygen species (ROS), and (poly) phenols (Miguel 2021a), which may affect regeneration efficiency (Seong et al. 2005; reviewed by Szechyńska-Hebda et al. 2007; Klimek-Szczykutowicz et al. 2022), could be mitigated by SA/salicylates. Ethylene biosynthesis decreased in mung bean (Lee et al. 1999) and in two barley cultivars under 300 mM NaCl stress (Torun et al. 2020), when exposed to SA. In cucumber seedlings, a SA treatment stimulated the antioxidant activity of the enzymes ascorbate peroxidase (APX), catalase (CAT), dehydroascorbate reductase (DHAR), guaiacol peroxidase (GPX), glutathione reductase (GR), and superoxide dismutase (SOD), decreased electrolyte leakage, H2O2 concentration, and thiobarbituric acid-reactive substances (TBARS), enhancing heat tolerance (Shi et al. 2006). In another study, SA inhibited CAT and APX activities while increasing SOD, peroxidase (POD), DHAR, and GR activities, decreased ROS levels and lipid peroxidation, and alleviating manganese toxicity in cucumber plants (Shi and Zhu 2008). Total polyphenol content in shaken cultures of watercress (*Nasturtium officinale* R. Br.) decreased with applied NaSA (Klimek-Szczykutowicz et al. 2022).

In some reports, the efficacy of applied SA/salicylates increased under stress conditions. For example, in *in vitro* propagation of three Cavendish banana (*Musa acuminata* Colla) cultivars, NaSA treatments under non-saline conditions were generally less effective than under salinity (120 mM NaCl) on the vegetative and physiological traits evaluated (Elazab and Youssef 2017).

In addition, other plant processes and attributes reported, in particular with NaSA application, may also influence regeneration efficiency, such as changes in photosynthetic pigment content (Elazab and Youssef 2017; Klimek-Szczykutowicz et al. 2022) and net photosynthetic and respiration rates (Al-Hakimi 2001), beneficial ion uptake and transport (Al-Hakimi and Hamada 2001; Elazab and Youssef 2017), enhanced growth (Al-Hakimi 2001; Elazab and Youssef 2017) and biomass parameters (Bhambhani et al. 2012; Klimek-Szczykutowicz et al. 2022), improved plant water relations (Al-Hakimi 2001), changes in secondary metabolite content (Silja and Satheeshkumar 2015; Klimek-Szczykutowicz et al. 2022), and so forth.

Differences between the effects of exogenously applied SA and NaSA on plants have been reported, such as on bud formation, growth parameters, heavy metal tolerance, plant immune response, total phenolic content, and antioxidant activity (Flores-Tena 1993; Zhang et al. 2001; Gondor et al. 2016; Karn et al. 2022). It is still unclear why SA and NaSA differ in their effects.

In the present investigation, the exogenous application of a specific concentration of NaSA (SA analog) showed a positive trend in *in vitro* adventitious organogenesis of a Spanish cucumber cultivar. This study can be extended to other cultivars and species. Further genetic, molecular, and physiological studies will help to understand, in particular, the role of this phenolic compound in *in vitro* plant regeneration.

## 5. Conclusions

From the present study, we conclude that *in vitro* shoot regeneration of a commercial cucumber cultivar was enhanced with the addition of sodium salicylate to the induction medium. These findings are an added value to address many biotechnological approaches in this economically important species, such as pathogen elimination, germplasm conservation, and large-scale multiplication.

## Abbreviations

BAP: 6-benzylaminopurine
IAA: Indole-3-acetic acid
KIN: Kinetin
MS: Murashige and Skoog (1962)

## Acknowledgements

The author is grateful to the Spanish Agency for International Development Cooperation (AECID) for the Ph.D. grant.

## References

Abrun A, Fattahi M, Hassani A, Avestan S (2016) Salicylic Acid and UV-B/C Radiation Effects on Growth and Physiological Traits of Satureja hortensis L. Not Sci Biol 8:170–175. https://doi.org/10.15835/nsb.8.2.9784

Akaike H (1973) Information theory and an extension of maximum likelihood principle. In: Proc. 2nd Int. Symp. on Information Theory. pp 267–281

Al-Hakimi AMA (2001) Alleviation of the adverse effects of NaCl on gas exchange and growth of wheat plants by ascorbic acid, thiamine and sodium salicylate. Pak J Biol Sci 4:762–765. https://doi.org/10.3923/pjbs.2001.762.765

Al-Hakimi Ama, Hamada AM (2001) Counteraction of salinity stress on wheat plants by grain soaking in ascorbic acid, thiamin or sodium salicylate. Biol Plant 44:253–261. https://doi.org/10.1023/a:1010255526903

Argueso CT, Ferreira FJ, Epple P, et al (2012) Two-Component Elements Mediate Interactions between Cytokinin and Salicylic Acid in Plant Immunity. PLOS Genet 8:e1002448. https://doi.org/10.1371/journal.pgen.1002448

Bakker PA, Ran L, Mercado-Blanco J (2014) Rhizobacterial salicylate production provokes headaches! Plant Soil 382:1–16. https://doi.org/10.1007/s11104-014-2102-0

Bergervoet JHW, Van der Mark F, Custers JBM (1989) Organogenesis versus embryogenesis from long-term suspension cultures of cucumber (Cucumis sativus L.). Plant Cell Rep 8:116–119. https://doi.org/10.1007/bf00716853

Bhambhani S, Karwasara VS, Dixit VK, Banerjee S (2012) Enhanced production of vasicine in Adhatoda vasica (L.) Nees. cell culture by elicitation. Acta Physiol Plant 34:1571–1578. https://doi.org/10.1007/s11738-011-0921-7

Bhutia KL, Meetei NT, Khanna VK (2018) In vitro direct regeneration of Dalle Khursani (Capsicum annum) from salicylic acid treated explants. J Pharmacogn Phytochem 7:1008–1012

Burza W, Malepszy S (1995) In vitro culture of Cucumis sativus L. XVIII. Plants from protoplasts through direct somatic embryogenesis. Plant Cell Tissue Organ Cult 41:259–266. https://doi.org/10.1007/BF00045090

Chee PP (1990) High Frequency of Somatic Embryogenesis and Recover of Fertile Cucumber Plants. HortScience 25:792–793. https://doi.org/10.21273/HORTSCI.25.7.792

Cleland CF, Tanaka O (1979) Effect of Daylength on the Ability of Salicylic Acid to Induce Flowering in the Long-day Plant Lemna gibba G3 and the Short-day Plant Lemna paucicostata 6746. Plant Physiol 64:421–424. https://doi.org/10.1104/pp.64.3.421

Damalas CA (2019) Improving drought tolerance in sweet basil (Ocimum basilicum) with salicylic acid. Sci Hortic 246:360–365. https://doi.org/10.1016/j.scienta.2018.11.005

Dong C-J, Liu X-Y, Xie L-L, et al (2020) Salicylic acid regulates adventitious root formation via competitive inhibition of the auxin conjugation enzyme CsGH3. 5 in cucumber hypocotyls. Planta 252:1–15. https://doi.org/10.1007/s00425-020-03467-2

Duman F, Aksoy A, Ozturk F, Erciyes AC (2010) Exogenous salicylate application affects the lead and copper accumulation characteristics of Lemna gibba L. Z Für Naturforschung C 65:675–680. https://doi.org/10.1515/znc-2010-11-1207

Elazab DS, Youssef M (2017) In vitro Response of Some Banana Cultivars to Salicylic Acid Treatment Under Salinity Stress. Assiut J Agric Sci 48:168–184. https://doi.org/10.21608/ajas.1999.5031

Fariduddin Q, Hayat S, Ahmad A (2003) Salicylic acid influences net photosynthetic rate, carboxylation efficiency, nitrate reductase activity, and seed yield in Brassica juncea. Photosynthetica 41:281–284. https://doi.org/10.1023/b:phot.0000011962.05991.6c

Flores-Tena J (1993) Efecto de tres salicilatos en el desarrollo de plantas de Solanum tuberosum cultivadas in vitro. Thesis, Universidad Nacional Autonoma de Mexico

Gondor OK, Pál M, Darkó É, et al (2016) Salicylic acid and sodium salicylate alleviate cadmium toxicity to different extents in maize (Zea mays L.). PLoS One 11:e0160157. https://doi.org/10.1371/journal.pone.0160157

Grozeva S, Velkov N (2014) In vitro plant regeneration of two cucumber (Cucumis sativum L.) genotypes: Effects of explant types and culture medium. Genetika 46:485–493. https://doi.org/10.2298/gensr1402485g

Harper JR, Balke NE (1981) Characterization of the inhibition of K+ absorption in oat roots by salicylic acid. Plant Physiol 68:1349–1353. https://doi.org/10.1104/pp.68.6.1349

Hayat Q, Hayat S, Irfan M, Ahmad A (2010) Effect of exogenous salicylic acid under changing environment: a review. Environ Exp Bot 68:14–25. https://doi.org/10.1016/j.envexpbot.2009.08.005

Hayat S, Siddiqui H, Damalas CA (eds) (2021) Salicylic Acid - A Versatile Plant Growth Regulator. Springer International Publishing. https://doi.org/10.1007/978-3-030-79229-9

Horváth E, Szalai G, Janda T (2007) Induction of abiotic stress tolerance by salicylic acid signaling. J Plant Growth Regul 26:290–300. https://doi.org/10.1007/s00344-007-9017-4

Jackman S (2020) pscl: Classes and Methods for R Developed in the Political Science Computational Laboratory. United States Studies Centre, University of Sydney, Sydney, New South Wales, Australia. R package version 1.5.5, https://github.com/atahk/pscl

Janda T, Gondor OK, Yordanova R, et al (2014) Salicylic acid and photosynthesis: signalling and effects. Acta Physiol Plant 36:2537–2546. https://doi.org/10.1007/s11738-014-1620-y

Jia S-R, Fu Y, Lin Y (1986) Embryogenesis and plant regeneration from cotyledon protoplast culture of cucumber (Cucumis sativus L.). J Plant Physiol 124:393–398. https://doi.org/10.1016/s0176-1617(86)80195-x

Jiang C-J, Shimono M, Sugano S, et al (2013) Cytokinins act synergistically with salicylic acid to activate defense gene expression in rice. Mol Plant Microbe Interact 26:287–296. https://doi.org/10.1094/mpmi-06-12-0152-r

Karn M, Sharma SK, Pal J, et al (2022) Efficacy of induced resistance chemicals in triggering defense mechanism of pomegranate plants against the pathogenic bacterium Xanthomonas axonopodis pv. punicae. Pharma Innov J 11:1656–1664

Khan MIR, Asgher M, Khan NA (2014) Alleviation of salt-induced photosynthesis and growth inhibition by salicylic acid involves glycinebetaine and ethylene in mungbean (Vigna radiata L.). Plant Physiol Biochem 80:67–74. https://doi.org/10.1016/j.plaphy.2014.03.026

Khan MIR, Iqbal N, Masood A, et al (2013) Salicylic acid alleviates adverse effects of heat stress on photosynthesis through changes in proline production and ethylene formation. Plant Signal Behav 8:e26374. https://doi.org/10.4161/psb.26374

Khan W, Prithiviraj B, Smith DL (2003) Photosynthetic responses of corn and soybean to foliar application of salicylates. J Plant Physiol 160:485–492. https://doi.org/10.1078/0176-1617-00865

Klessig DF, Tian M, Choi HW (2016) Multiple Targets of Salicylic Acid and Its Derivatives in Plants and Animals. Front Immunol 7:1–10. https://doi.org/10.3389/fimmu.2016.00206

Klimek-Szczykutowicz M, Dziurka M, Blažević I, et al (2022) Impacts of elicitors on metabolite production and on antioxidant potential and tyrosinase inhibition in watercress microshoot cultures. Appl Microbiol Biotechnol 106:619–633. https://doi.org/10.1007/s00253-021-11743-8

Koo YM, Heo AY, Choi HW (2020) Salicylic acid as a safe plant protector and growth regulator. Plant Pathol J 36:1–10. https://doi.org/10.5423/ppj.rw.12.2019.0295

Kračun-Kolarević M, Dmitrović S, Filipović B, et al (2015) Influence of sodium salicylate on rosmarinic acid, carnosol and carnosic acid accumulation by Salvia officinalis L. shoots grown in vitro. Biotechnol Lett 37:1693–1701. https://doi.org/10.1007/s10529-015-1825-1

Kreuger M, Meer W van der, Postma E, et al (1996) Genetically stable cell lines of cucumber for the large-scale production of diploid somatic embryos. Physiol Plant 97:303–310. https://doi.org/10.1034/j.1399-3054.1996.970213.x

Latif F, Ullah F, Mehmood S, et al (2016) Effects of salicylic acid on growth and accumulation of phenolics in Zea mays L. under drought stress. Acta Agric Scand Sect B—Soil Plant Sci 66:325–332. https://doi.org/10.1080/09064710.2015.1117133

Lee J-H, Jin ES, Kim WT (1999) Inhibition of auxin-induced ethylene production by salicylic acid in mung bean hypocotyls. J Plant Biol 42:1–7. https://doi.org/10.1007/BF03031140

Lee S, Kim S-G, Park C-M (2010) Salicylic acid promotes seed germination under high salinity by modulating antioxidant activity in Arabidopsis. New Phytol 188:626–637. https://doi.org/10.1111/j.1469-8137.2010.03378.x

Li XM, Ma LJ, Bu N, et al (2014) Effects of salicylic acid pre-treatment on cadmium and/or UV-B stress in soybean seedlings. Biol Plant 58:195–199. https://doi.org/10.1007/s10535-013-0375-4

Liao D, Chen X, Chen A, et al (2015) The Characterization of Six Auxin-Induced Tomato GH3 Genes Uncovers a Member, SlGH3.4, Strongly Responsive to Arbuscular Mycorrhizal Symbiosis. Plant Cell Physiol 56:674–687. https://doi.org/10.1093/pcp/pcu212

Lou H, Kako S (1994) Somatic Embryogenesis and Plant Regeneration in Cucumber. HortScience 29:906–909. https://doi.org/10.21273/HORTSCI.29.8.906

Luo J-P, Jiang S-T, Pan L-J (2001) Enhanced somatic embryogenesis by salicylic acid of Astragalus adsurgens Pall.: relationship with H2O2 production and H2O2-metabolizing enzyme activities. Plant Sci 161:125–132. https://doi.org/10.1016/S0168-9452(01)00401-0

Madany MMY, Obaid WA, Hozien W, et al (2020) Salicylic acid confers resistance against broomrape in tomato through modulation of C and N metabolism. Plant Physiol Biochem 147:322–335. https://doi.org/10.1016/j.plaphy.2019.12.028

Malabadi RB, Teixeira da Silva JA, Nataraja K (2008) Salicylic acid induces somatic embryogenesis from mature trees of Pinus roxburghii (Chir pine) using TCL technology. Tree For Sci Biotechnol 2:34–39

Malepszy S, Nadolska-Orczyk A (1983) In vitro Culture of Cucumis sativus I. Regeneration of Plantlets From Callus Formed by Leaf Expiants. Z Für Pflanzenphysiol 111:273–276. https://doi.org/10.1016/S0044-328X(83)80086-5

Malepszy S, Solarek E (1986) In vitro culture of Cucumis sativus L. IV. Conditions for cell suspension. Genet Pol 27:249–253

Mann KK, Schmann AW, Spann TM (2011) Response of citrus to exogenously applied salicylate compounds during abiotic and biotic stress. In: Proceedings of the Florida State Horticultural Society. pp 101–110

Maurya B, Rai KK, Pandey N, et al (2019) Influence of salicylic acid elicitation on secondary metabolites and biomass production in in-vitro cultured Withania coagulans (L.) Dunal. Plant Arch 19:1308–1316

Miguel JF (2017) Studies on regeneration and genetic transformation in cucumber (Cucumis sativus L.) via Agrobacterium tumefaciens (in Spanish). PhD Thesis. Universidad Politécnica de Valencia. https://doi.org/10.4995/thesis/10251/90405

Miguel JF (2021a) Influence of High Concentrations of Copper Sulfate on In Vitro Adventitious Organogenesis of Cucumis sativus L. bioRxiv 2021.02.24.432794. https://doi.org/10.1101/2021.02.24.432794

Miguel JF (2021b) Effect of Light Conditions on In Vitro Adventitious Organogenesis of Cucumber Cultivars. 2021.12.13.472500. https://doi.org/10.1101/2021.12.13.472500

Mohamed HI, El-Shazly HH, Badr A (2020) Role of salicylic acid in biotic and abiotic stress tolerance in plants. In: Plant Phenolics in Sustainable Agriculture. Springer, pp 533–554. https://doi.org/10.1007/978-981-15-4890-1_23

Nandy S, Das T, Dey A (2021) Role of Jasmonic Acid and Salicylic Acid Signaling in Secondary Metabolite Production. Jasmonates Salicylates Signal Plants 87–113. https://doi.org/10.1007/978-3-030-75805-9_5

Pacheco G, Cardoso SRS, Gagliardi RF, et al (2008) Genetic and epigenetic analyses of in vitro-grown plants of Arachis villosulicarpa Hoehne (Leguminosae) obtained from seed explants through different regeneration pathways. J Hortic Sci Biotechnol 83:737–742. https://doi.org/10.1080/14620316.2008.11512453

Palmer IA, Chen H, Chen J, et al (2019) Novel Salicylic Acid Analogs Induce a Potent Defense Response in Arabidopsis. Int J Mol Sci 20:3356. https://doi.org/10.3390/ijms20133356

Pérez-Llorca M, Muñoz P, Müller M, Munné-Bosch S (2019) Biosynthesis, Metabolism and Function of Auxin, Salicylic Acid and Melatonin in Climacteric and Non-climacteric Fruits. Front Plant Sci 10:136. https://doi.org/10.3389/fpls.2019.00136

Pokotylo I, Kravets V, Ruelland E (2019) Salicylic Acid Binding Proteins (SABPs): The Hidden Forefront of Salicylic Acid Signalling. Int J Mol Sci 20:E4377. https://doi.org/10.3390/ijms20184377

PubChem (2022a) Sodium salicylate. https://pubchem.ncbi.nlm.nih.gov/compound/16760658. Accessed 10 Sep 2022

PubChem (2022b) Salicylic acid. https://pubchem.ncbi.nlm.nih.gov/compound/338. Accessed 10 Sep 2022

Punja ZK, Abbas N, Sarmento GG, Tang FA (1990) Regeneration of Cucumis sativus var. sativus and C. sativus var. hardwickii, C. melo, and C. metuliferus from explants through somatic embryogenesis and organogenesis. Plant Cell Tissue Organ Cult 21:93–102. https://doi.org/10.1007/BF00033427

Quiroz-Figueroa F, Méndez-Zeel M, Larqué-Saavedra A, Loyola-Vargas V (2001) Picomolar concentrations of salicylates induce cellular growth and enhance somatic embryogenesis in Coffea arabica tissue culture. Plant Cell Rep 20:679–684. https://doi.org/10.1007/s002990100386

R Core Team (2021) R: A language and environment for statistical computing. R Foundation for Statistical Computing, Vienna. https://www.R-project.org

Raskin I, Skubatz H, Tang W, Meeuse BJ (1990) Salicylic acid levels in thermogenic and non-thermogenic plants. Ann Bot 66:369–373. https://doi.org/10.1093/oxfordjournals.aob.a088037

Rhoads DM, McIntosh L (1992) Salicylic acid regulation of respiration in higher plants: alternative oxidase expression. Plant Cell 4:1131–1139. https://doi.org/10.2307/3869481

Rivas-San Vicente M, Plasencia J (2011) Salicylic acid beyond defence: its role in plant growth and development. J Exp Bot 62:3321–3338. https://doi.org/10.1093/jxb/err031

Sakhanokho HF, Kelley RY (2009) Influence of salicylic acid on in vitro propagation and salt tolerance in Hibiscus acetosella and Hibiscus moscheutos (cv ‘Luna Red’). Afr J Biotechnol 8:1474–1481.

Saleem M, Fariduddin Q, Janda T (2020) Multifaceted Role of Salicylic Acid in Combating Cold Stress in Plants: A Review. J Plant Growth Regul 40:464–485. https://doi.org/10.1007/s00344-020-10152-x

Schwarz G (1978) Estimating the Dimension of a Model. Ann Stat 6:461–464. https://doi.org/10.1214/aos/1176344136

Sellers K, Lotze T, Raim A, Raim MA (2019) Package ‘COMPoissonReg’. https://github.com/lotze/COMPoissonReg

Selvaraj N, Vasudevan A, Manickavasagam M, et al (2007) High frequency shoot regeneration from cotyledon explants of cucumber via organogenesis. Sci Hortic 112:2–8. https://doi.org/10.1016/j.scienta.2006.12.037

Seo S-H, Bai D-G, Park H-Y (2000) High frequency shoot regeneration from leaf explants of cucumber. J Plant Biotechnol 2:51–54.

Seong ES, Song KJ, Jegal S, et al (2005) Silver nitrate and aminoethoxyvinylglycine affect Agrobacterium-mediated apple transformation. Plant Growth Regul 45:75–82. https://doi.org/10.1007/s10725-004-6126-y

Shakirova FM, Sakhabutdinova AR, Bezrukova MV, et al (2003) Changes in the hormonal status of wheat seedlings induced by salicylic acid and salinity. Plant Sci 164:317–322. https://doi.org/10.1016/s0168-9452(02)00415-6

Sharma A, Sidhu GPS, Araniti F, et al (2020) The role of salicylic acid in plants exposed to heavy metals. Molecules 25:540. https://doi.org/10.3390/molecules25030540

Kosakivska IV, Shcherbatiuk MM, Voytenko LV (2020) Profiling of hormones in plant tissues: history, modern approaches, use in biotechnology. Biotechnologia Acta 13:14–25. https://doi.org/10.15407/biotech13.04.014

Shemi R, Wang R, Gheith E-SMS, et al (2021) Effects of salicylic acid, zinc and glycine betaine on morpho-physiological growth and yield of maize under drought stress. Sci Rep 11:3195. https://doi.org/10.1038/s41598-021-82264-7

Shettel NL, Balke NE (1983) Plant growth response to several allelopathic chemicals. Weed Sci 31:293–298. https://doi.org/10.1017/s0043174500069034

Shetty K, Shetty GA, Nakazaki Y, et al (1992) Stimulation of benzyladenine-induced in vitro shoot organogenesis in Cucumis melo L. by proline, salicylic acid and aspirin. Plant Sci 84:193–199. https://doi.org/10.1016/0168-9452(92)90134-8

Shi Q, Bao Z, Zhu Z, et al (2006) Effects of different treatments of salicylic acid on heat tolerance, chlorophyll fluorescence, and antioxidant enzyme activity in seedlings of Cucumis sativa L. Plant Growth Regul 48:127–135. https://doi.org/10.1007/s10725-005-5482-6

Shi Q, Zhu Z (2008) Effects of exogenous salicylic acid on manganese toxicity, element contents and antioxidative system in cucumber. Environ Exp Bot 63:317–326. https://doi.org/10.1016/j.envexpbot.2007.11.003

Silja PK, Satheeshkumar K (2015) Establishment of adventitious root cultures from leaf explants of Plumbago rosea and enhanced plumbagin production through elicitation. Ind Crops Prod 76:479–486. https://doi.org/10.1016/j.indcrop.2015.07.021

Sohag AAM, Tahjib-Ul-Arif M, Brestic M, et al (2020) Exogenous salicylic acid and hydrogen peroxide attenuate drought stress in rice. Plant Soil Environ 66:7–13. https://doi.org/10.17221/472/2019-pse

Su YH, Zhang XS (2014) Chapter Two - The Hormonal Control of Regeneration in Plants. In: Galliot B (ed) Current Topics in Developmental Biology. Academic Press, pp 35–69. https://doi.org/10.1016/b978-0-12-391498-9.00010-3

Szechyńska-Hebda M, Skrzypek E, Dąbrowska G, et al (2007) The role of oxidative stress induced by growth regulators in the regeneration process of wheat. Acta Physiol Plant 29:327–337. https://doi.org/10.1007/s11738-007-0042-5

Torun H, Novák O, Mikulík J, et al (2020) Timing-dependent effects of salicylic acid treatment on phytohormonal changes, ROS regulation, and antioxidant defense in salinized barley (Hordeum vulgare L.). Sci Rep 10:13886. https://doi.org/10.1038/s41598-020-70807-3

Trulson AJ, Shahin EA (1986) In vitro plant regeneration in the genus Cucumis. Plant Sci 47:35–43. https://doi.org/10.1016/0168-9452(86)90008-7

Uthpala TGG, Marapana R, Lakmini K, Wettimuny DC (2020) Nutritional bioactive compounds and health benefits of fresh and processed cucumber (Cucumis sativus L.). Sumerianz J Biotechnol 3:75–82. https://doi.org/10.13140/RG.2.2.17510.04161

Uzodike EB, Onuoha IN (2009) The effect of cucumber (Cucumbis savitus) extract on acid induced corneal burn in guinea pigs. J Niger Optom Assoc 15:3–7. https://doi.org/10.4314/jnoa.v15i1.55592

Vasudevan A, Ganapathi A, Choi CW (2006) Effect of ethylene inhibitors on in vitro shoot multiplication and their impact on ethylene production in cucumber (Cucumis sativus L.). J Plant Biotechnol 33:249–255. https://doi.org/10.5010/jpb.2006.33.4.249

Wang S, Ku SS, Ye X, et al (2015) Current status of genetic transformation technology developed in cucumber (Cucumis sativus L.). J Integr Agric 14:469–482. https://doi.org/10.1016/S2095-3119(14)60899-6

Wehner TC (1981) In vitro adventitious shoot and root formation of cultivars and lines of Cucumis sativus L. HortScience 16:759–760

Wu J, Liu S, Guan X, et al (2014) Genome-wide identification and transcriptional profiling analysis of auxin response-related gene families in cucumber. BMC Res Notes 7:1–13. https://doi.org/10.1186/1756-0500-7-218

Zhang S, Reddy MS, Kokalis-Burelle N, et al (2001) Lack of induced systemic resistance in peanut to late leaf spot disease by plant growth-promoting rhizobacteria and chemical elicitors. Plant Dis 85:879–884. https://doi.org/10.1094/pdis.2001.85.8.879

Zhong Q, Hu H, Fan B, et al (2021) Biosynthesis and Roles of Salicylic Acid in Balancing Stress Response and Growth in Plants. Int J Mol Sci 22:11672. https://doi.org/10.3390/ijms222111672

Ziv M, Gadasi G (1986) Enhanced embryogenesis and plant regeneration from cucumber (Cucumis sativus L.) callus by activated charcoal in solid/liquid double-layer cultures. Plant Sci 47:115–122. https://doi.org/10.1016/0168-9452(86)90058-0

